# Evolutionary footprints reveal insights into plant microRNA biogenesis

**DOI:** 10.1101/129684

**Authors:** Uciel Chorostecki, Belen Moro, Arantxa M.L. Rojas, Juan M. Debernardi, Arnaldo L. Schapire, Cedric Notredame, Javier F. Palatnik

## Abstract

MicroRNAs (miRNAs) are endogenous small RNAs that recognize target sequences by base complementarity. They are processed from longer precursors that harbor a fold-back structure. Plant miRNA precursors are quite variable in size and shape, and are recognized by the processing machinery in different ways. However, ancient miRNAs and their binding sites in target genes are conserved during evolution. Here, we designed a strategy to systematically analyze *MIRNAs* from different species generating a graphical representation of the conservation of the primary sequence and secondary structure. We found that plant *MIRNAs* have evolutionary footprints that go beyond the small RNA sequence itself, yet, their location along the precursor depends on the specific *MIRNA*. We show that these conserved regions correspond to structural determinants recognized during the biogenesis of plant miRNAs. Furthermore, we found that the members of the miR166 family have unusual conservation patterns and demonstrated that the recognition of these precursors in vivo differs from other known miRNAs. Our results describe a link between the evolutionary conservation of plant *MIRNAs* and the mechanisms underlying the biogenesis of these small RNAs, and show that the *MIRNA* pattern of conservation can be used to infer the mode of miRNA biogenesis.

## Introduction

MiRNAs are small RNAs of 20-22 nt that originate from endogenous loci and regulate other target RNAs by base complementarity in animals and plants (Rogers and Chen, 2013; Bologna and Voinnet, 2014). They have emerged and specialized independently in both kingdoms, which likely explains differences in their biogenesis and action modes (Axtell et al., 2011; Cui et al., 2016). MiRNAs are transcribed as longer precursors harboring an imperfect fold-back structure, with the small RNA embedded in one of its arms. These precursors contain spatial cues that are recognized during the biogenesis of the small RNAs (Bologna and Voinnet, 2014; Ha and Kim, 2014).

A typical animal miRNA primary transcript harbors a fold-back structure that consists of a ^∼^35 bp stem and a terminal loop that are flanked by single-stranded RNA (ssRNA) segments (Ha and Kim, 2014). These transcripts are processed by the microprocessor, a complex that contains the RNAse type III Drosha, which recognizes the transition of the ssRNA and the double-stranded region (dsRNA) of the stem loop, and produces a first cut ^∼^11 bp away of this ssRNA-dsRNA junction (Ha and Kim, 2014). The resulting pre-miRNA is exported to the cytoplasm where Dicer performs the second cut by ^∼^22 nt away from the first cleavage site, releasing a miRNA/miRNA* duplex (Ha and Kim, 2014). The miRNA is finally incorporated into an AROGNAUTE (AGO) complex, which is responsible for the activity of the small RNA (Axtell et al., 2011; Bologna et al., 2013a; Bologna and Voinnet, 2014; Ha and Kim, 2014).

Plant miRNA precursors are much more variable in size and shape than their animal counterparts, and they are completely processed in the nucleus by a complex harboring DICER-LIKE1 (DCL1) (Axtell et al., 2011; Rogers and Chen, 2013; Bologna and Voinnet, 2014). That plant miRNAs can be processed in different ways likely explains the lack of features common to all precursors (Bologna et al., 2013b). Rather, plant miRNA precursors can be classified into several groups. One group harbors plant miRNA precursors with a ∼15-17 nt stem below the miRNA/miRNA* (lower stem), which specifies the position of the first cut by DCL1 (Mateos et al., 2010; Song et al., 2010; Werner et al., 2010; Bologna et al., 2013b; Zhu et al., 2013). A second cut by DCL1, ^∼^21 nt away from the first cleavage site releases the miRNA/miRNA*. These precursors are processed in a base-to-loop direction resembling the processing of animal miRNAs.

However, another group of plant miRNA precursors are processed by a first cleavage below the terminal loop and from there processing continues towards the base of the precursor (Addo-Quaye et al., 2009; Bologna et al., 2009; Bologna et al., 2013b). These loop-to-base processed precursors have a structured dsRNA region above of the miRNA/miRNA* (upper stem), which is recognized by the processing machinery (Addo-Quaye et al., 2009; Bologna et al., 2009; Bologna et al., 2013b; Kim et al., 2016). In addition, some plant miRNA precursors are processed sequentially by several cuts instead of the usual two found in animals (Kurihara and Watanabe, 2004; Addo-Quaye et al., 2009; Bologna et al., 2009; Zhang et al., 2010; Bologna et al., 2013b).

Since the discovery of plant *MIRNAs*, it has been pointed out that conservation in distant species is only clearly seen in the miRNA/miRNA* region (Reinhart et al., 2002). The conservation of the actual miRNA sequences can be readily explained by the conserved recognition sites of cognate target genes (Reinhart et al., 2002; Allen et al., 2004; Jones-Rhoades and Bartel, 2004), a characteristic that has been exploited to predict miRNA target genes (Jones-Rhoades and Bartel, 2004; Chorostecki et al., 2012). However, the precursor of the ancient miR319 harbors a second conserved region in the precursor stem above the miRNA/miRNA* (Palatnik et al., 2003; Axtell and Bartel, 2005; Warthmann et al., 2008; Addo-Quaye et al., 2009; Bologna et al., 2009; Li et al., 2011; Sobkowiak et al., 2012), showing that additional conserved sequences exist in at least certain *MIRNAs*.

Here, we performed a global analysis of *MIRNA* sequences in different plant species. We designed a graphic representation that displays quantitative information on the conservation of the primary sequences and secondary structures. We found evolutionary footprints in plant *MIRNAs* that go beyond the miRNA/miARN* region and reveal conservation of miRNA processing. Precursors processed in a loop-to-base or base-to-loop direction, by two or more cuts have all distinct evolutionary footprints, suggesting that the miRNA processing pathway can be inferred from the conservation pattern of a *MIRNA*. As a proof of principle, we used this approach to identify new miRNA processing determinants and found that the evolutionarily conserved miR166 miRNAs require just a few bases outside the miR166/miR166* region for their biogenesis, demonstrating that their precursor recognition differs from other known miRNAs. The results describe a strong link between the evolutionary conservation of plant *MIRNAs* and the mechanisms underlying the biogenesis of the small RNAs.

## Results and discussion

### Identification of plant miRNA precursors in different species

*MIRNAs* which encode similar or identical small RNAs are usually grouped into a single family (Meyers et al., 2008). There are 29 families of miRNAs conserved at least in dicotyledonous plants (Cuperus et al., 2011; Chavez Montes et al., 2014), which are represented by 96 different precursors in *Arabidopsis thaliana* (miRBASE, release 21) (Figure 1), although the exact number might vary depending on whether miR156/157, miR165/166, miR170/171 or miR159/miR319 are considered part of a single family or superfamily (Meyers et al., 2008; Cuperus et al., 2011). Considering that miRNA precursors of the same family can be processed in different ways (Bologna et al., 2013b), we analyzed the conservation of orthologous *MIRNAs*, instead of grouping all different members of each miRNA family.

**Figure 1:**
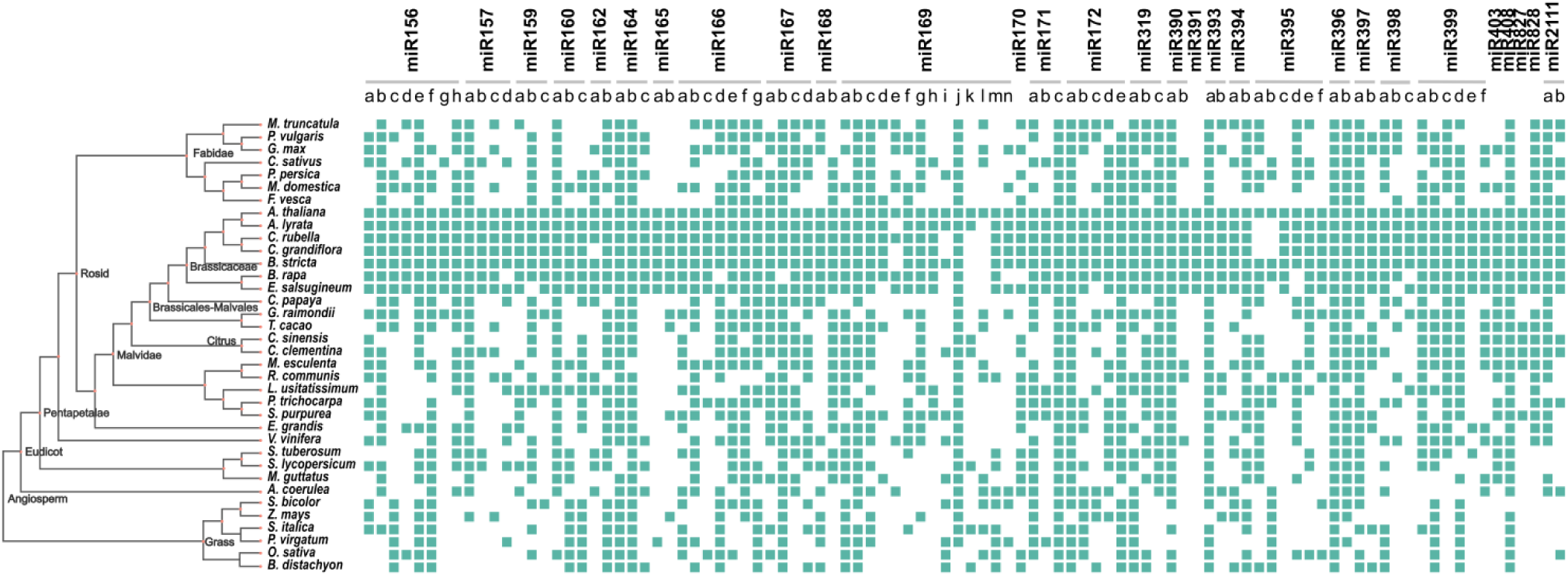
Identification of Arabidopsis *MIRNA* orthologues in dicotyledonous and monocotyledonous plants. Representation of putative orthologs detected in dicotyledonous and monocotyledonous species for 96 *MIRNAs*.

Reciprocal BLAST was used to identify putative orthologous genes to the Arabidopsis miRNAs in the genomes of 30 dicotyledonous and 6 monocotyledonous species available in the Phytozome database (version 11) (Figure 1). Starting with the 96 Arabidopsis *MIRNAs*, we identified 2112 putative orthologous sequences in other species (Figure 1, Supplementary Dataset 1). This large group of sequences will not cover exhaustively all miRNA precursors corresponding to the conserved miRNA families in the 36 angiosperms analyzed, but it should provide enough sequence information to allow a general analysis of their sequence conservation.

### Visualization of *MIRNA* primary sequence and secondary structure of different species

The putative orthologous sequences of each Arabidopsis conserved *MIRNA* were used to perform a multiple alignment using T-coffee (Chang et al., 2014), and 96 different alignments were generated. We analyzed separately dicots alone (1886 precursors, Supplementary Figure 1, Figure 2A) or dicots together with monocots (2112 precursors, Supplementary Figure 2). The secondary structure of each *MIRNA* sequence was also predicted using RNA fold (Lorenz et al., 2011) (Supplementary Figure 3). To visualize the complex data obtained we generated a representation of the miRNA precursors based on Circos (Krzywinski et al., 2009) (Supplementary Figure 4 and 5, Figure 2B-G). In this representation, the inner ring shows a histogram of the frequency and distribution of paired (Figure 2B-G, green bars) and unpaired bases (Figure 2B-G, purple bars) for every position in the precursor, which, therefore, quantitatively indicates the conservation of secondary structures in different species. At the same time, the outer Circos data shows the nucleotide sequence of the Arabidopsis precursor maintaining the color conservation of the multiple sequence alignment consensus (Supplementary Figure 4 and 5, Figure 2B-G). The Circos-based studies were performed in dicots (Figure 2B-D, Supplementary Figure 4) and dicots together with monocots (Figure 2E-G, Supplementary Figure 5), as we had done for the T-coffee alignments (Supplementary Figure 1 and 2).

**Figure 2:**
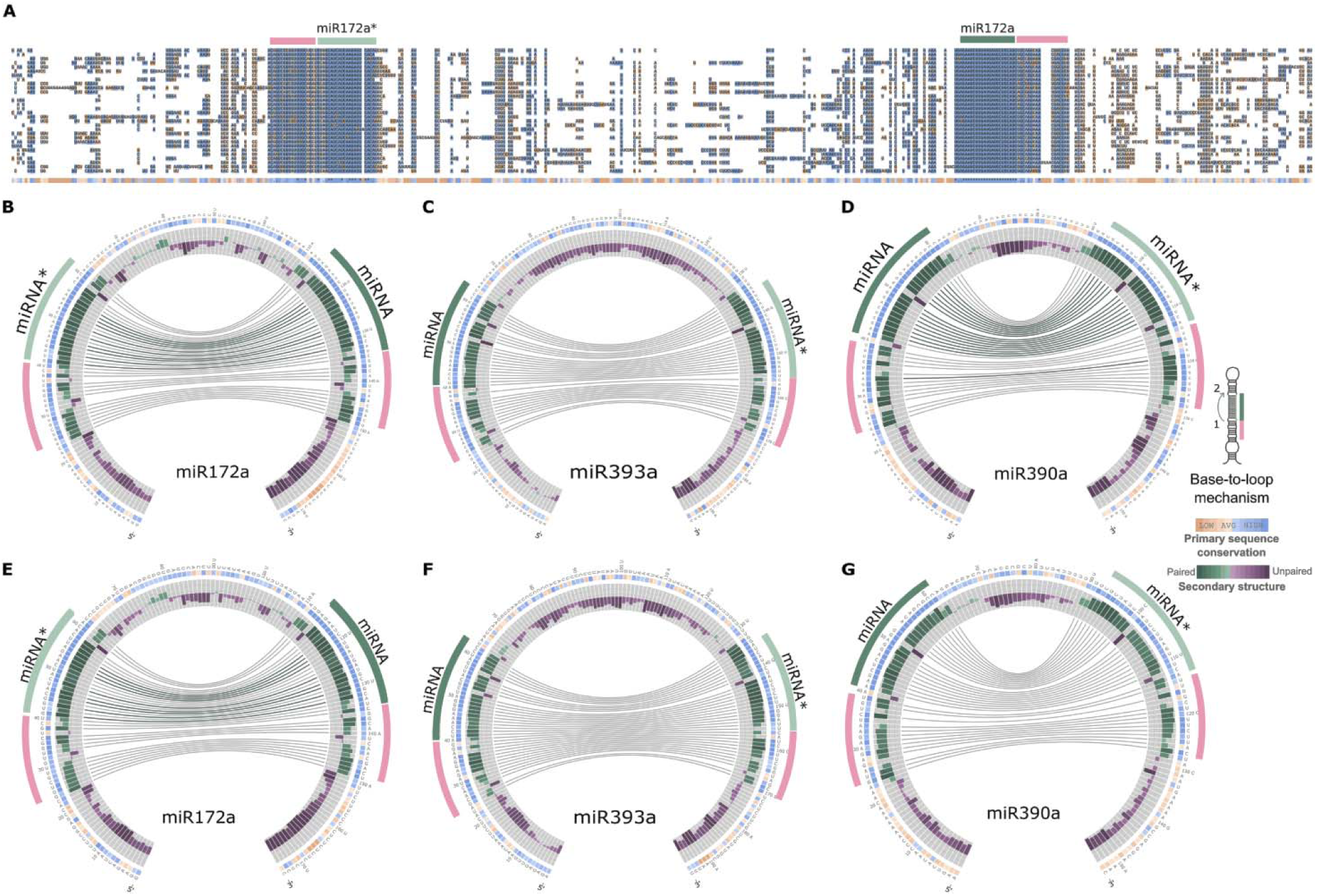
Circos representation of miRNA precursors processed in a base-to-loop direction. (A) Alignment of miR172a precursors from *A. lyrata* (top)*, M. domestica, M. truncatula, S. lycopersicum, B. rapaFPsc, E. grandis, C. grandiflora, P. persica, C. sinensis, L. usitatissimum, C. clementina, G. max, V. vinifera, R. communis, S. purpurea, B. stricta, C. sativus, A. coerulea, M. guttatus, M. esculenta, E. salsugineum, C. papaya, A. thaliana, C. rubella, T. cacao, P. trichocarpa, P. vulgaris, G. raimondii, F. vesca and S. tuberosum* (bottom). (B-G) Circos representation of miR172a (B, E), miR393a (C, F) and miR390a (D, G) precursors in dicots (B-D) and dicots and monocots (E-G). Conserved sequences are indicated with the same color code as the alignment (A). Green bars indicate bases that tend to form dsRNA regions which are quantitatively indicated by the height of the bars. Connecting lines refer to bases that are interacting in the secondary structure of the precursors, green lines refer to bases that interact 100%, while gray lines show bases interacting in at least 50% of the species. Purple bars refer to bases that tend to be ssRNA regions. The miRNA is indicated with green line, while the miRNA* is light green. A conserved region that corresponds to a ^∼^15nt lower stem is indicated with a pink line. The reference sequence is the Arabidopsis miRNA precursor. Note that sequences below the lower stem and the loop a mostly unpaired (purple bars). The inset (right) shows a scheme of a precursor processed by a base-to-loop mechanism.

**Figure 3:**
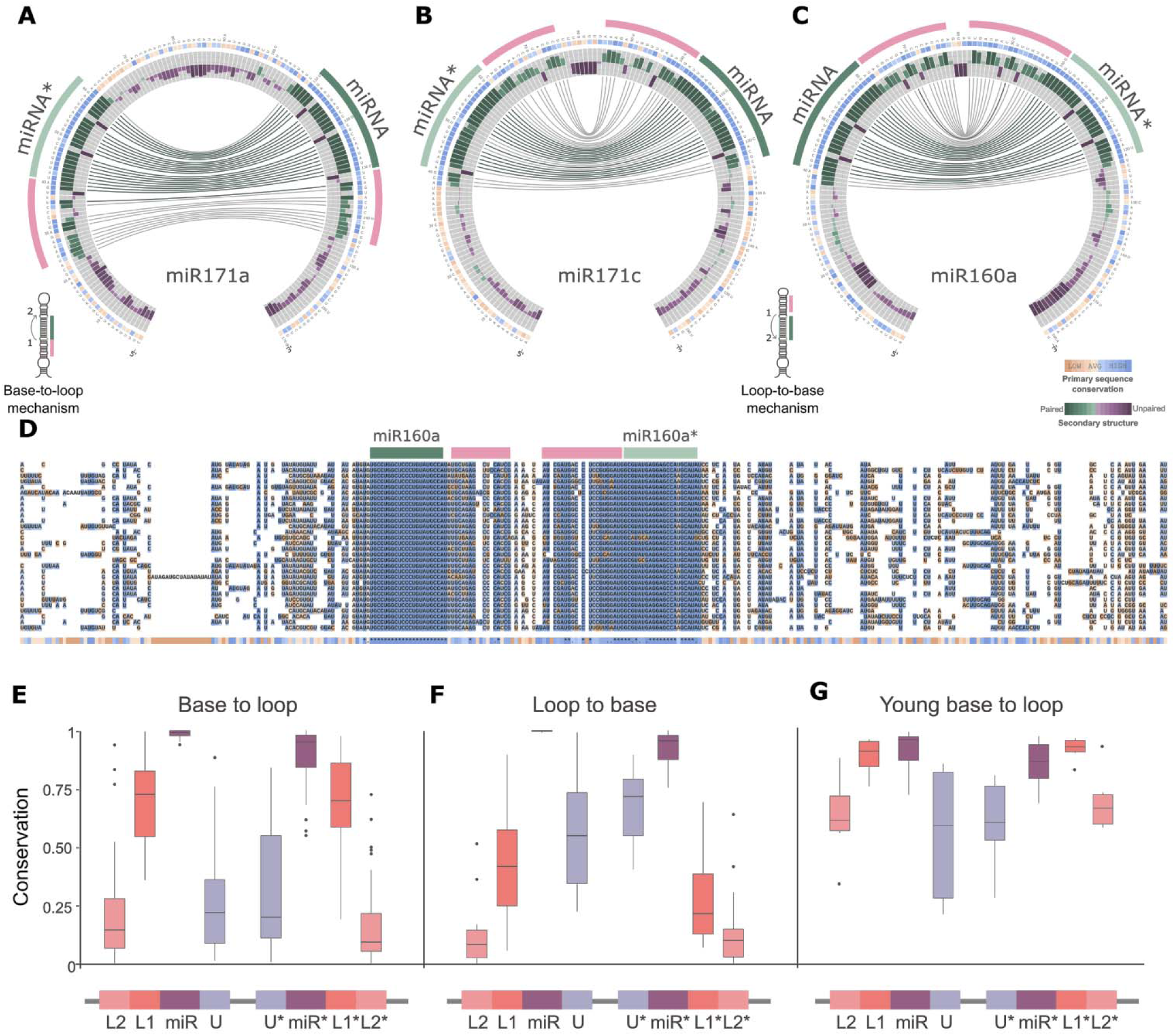
Conservation and divergence of precursors processed in different directions. (A-C) Circos representation of miR171a (A), miR171c (B) and miR160a (C). miR171a is processed from the base, while miR171c and miR160a are processed from the loop. Note the different position of the additional conserved regions (pink line) in the precursors according their processing direction. The insets show schemes of precursors processed in base-to-loop or a loop-to-base direction. (D) Alignment of miR171c precursors from *A. lyrata* (top), *M. domestica, M. truncatula, S. lycopersicum, B. rapaFPsc, E. grandis, C. grandiflora, P. persica, C. sinensis, L. usitatissimum, C. clementina, G. max, V. vinifera, R. communis, S. purpurea, B. stricta, C. sativus, A. coerulea, M. guttatus, M. esculenta, E. salsugineum, C. papaya, A. thaliana, C. rubella, T. cacao, P. trichocarpa, P. vulgaris, G. raimondii, F. vesca and S. tuberosum* (bottom). (E-G) Box plot showing the conservation of different precursor regions using phastCons for precursors processed in base-to-loop (E) or loop-to-base (F) direction. (G) Analysis using young *MIRNAs* processed in base-to-loop direction. The band inside the box represents the median, the bottom and top of the box are the first (Q1) and third (Q3) quartiles, dots are outliers, upper whisker denotes min (max(x), Q3 + 1.5 * (Q3 - Q1)) and lower whisker denotes max(min(x), Q1 - 1.5 * (Q3 - Q1)).

**Figure 4:**
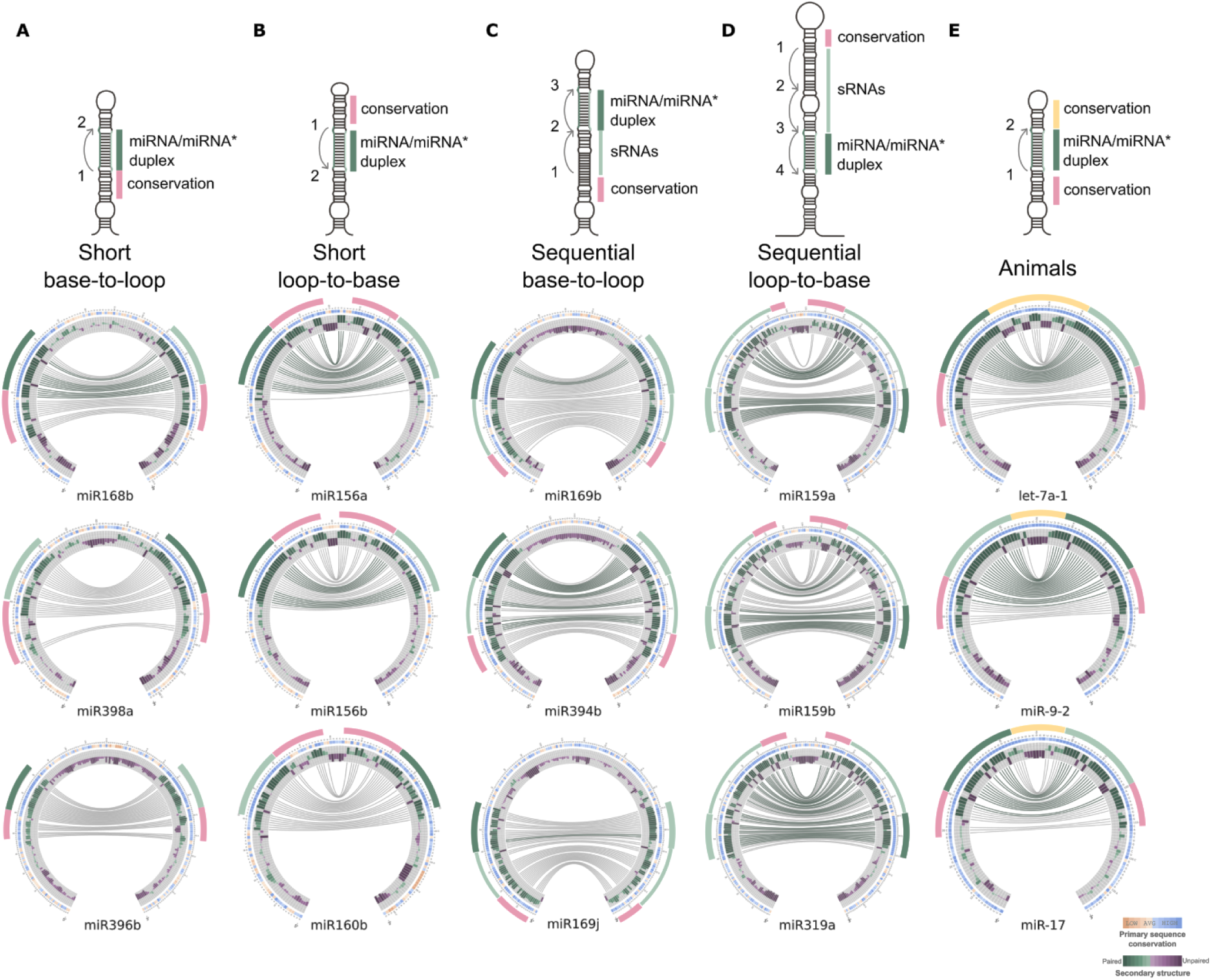
miRNA biogenesis pathways tune *MIRNA* conservation during evolution. (A-D) Circos representation of precursors processing through different directions: base to loop (A), loop to base (B), and sequentially processed precursors from the base (C) or the loop (D). The miRNA is indicated by a green line, while the miRNA* is light green. Other small RNAs are indicated by a narrow green line. A conserved region that corresponds to a lower or upper stem is indicated by a pink line. The conservation of the distal region in animals is indicated in yellow. Top, schemes representing the different processing pathways. (E) Circos representation of animal *MIRNAs* (analyzed species include *Bos taurus, Canis familiaris, Equus caballus, Gallus gallus, Gorilla gorilla, Homo sapiens, Macaca mulatta, Monodelphis domestica, Mus musculus, Ornithorhynchus anat nus, Petromyzon marinus, Sus scrofa and Xenopus tropicalis*).

**Figure 5:**
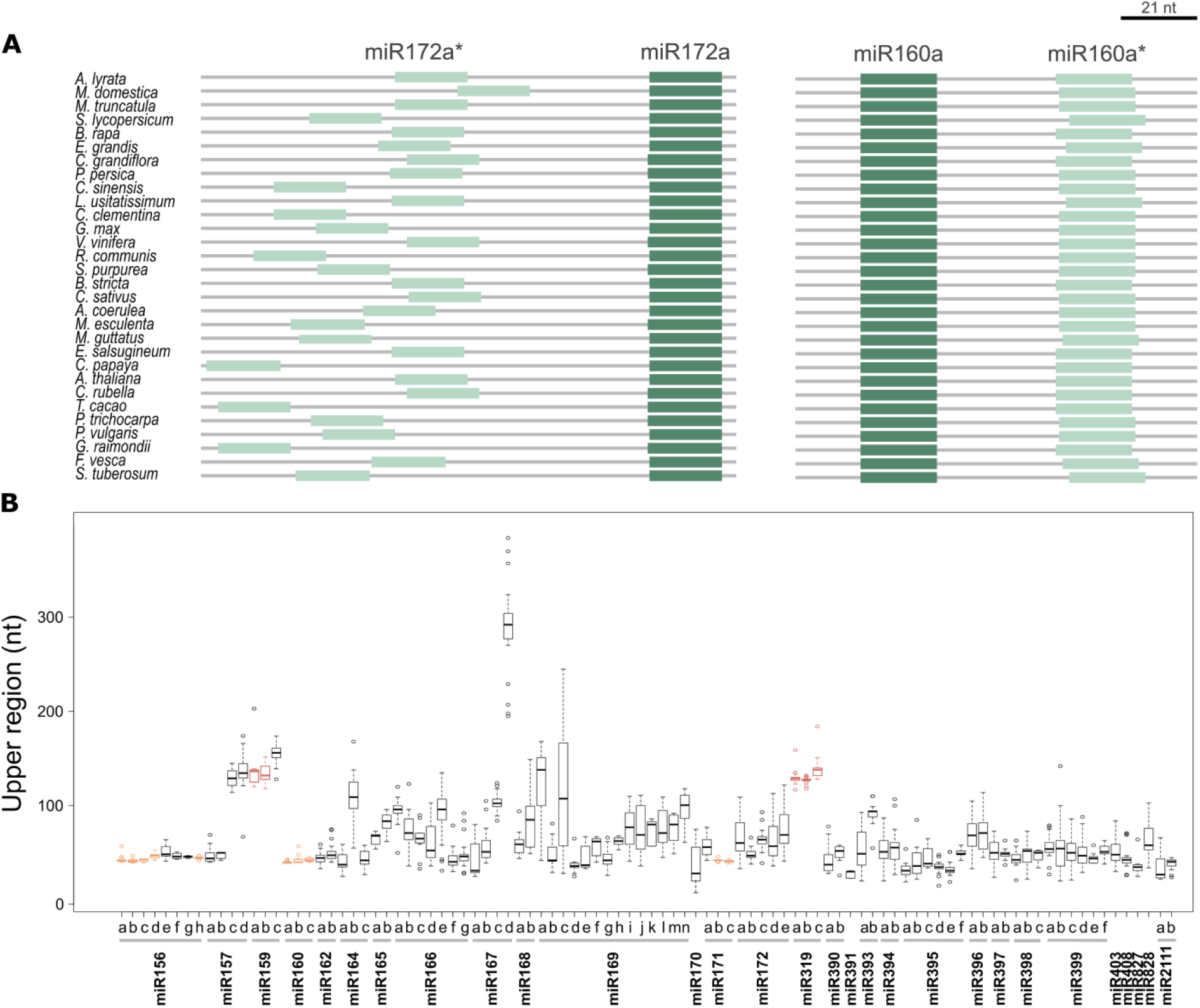
Precursor length divergence in different species. (A) Schemes showing the relative positions of the miR172a and miR172a*, and miR160a and miR160* in the precursor sequences of different species. Note the changes in the relative positions for miR172*, and the conservation for miR160*. (B) Box plot showing the length (nt) between the miRNA and the miRNA* of conserved precursors. Short loop-to-base precursors experimentally validated are shown in orange and sequential loop-to-base precursors are shown in red. The band inside the box represents the median, the bottom and top of the box are the first (Q1) and third quartiles (Q3), dots are outliers, upper whisker denotes min(max(x), Q3 + 1.5 * (Q3 - Q1)) and lower whisker denotes max(min(x), Q1 - 1.5 * (Q3 - Q1)).

An inspection of the *MIRNA* alignments revealed that the plant precursors have evolutionary footprints in addition to the conservation of the miRNA/miRNA* region, however, the length and relative position of the conserved regions varied among the different *MIRNAs* (Supplementary Figure 1 and 2). We looked in more detail into the alignment of *MIR172a* (Figure 2A), whose precursor structure-function relationship has already been experimentally studied in detail (Mateos et al., 2010; Werner et al., 2010). In this case, the miR172/miR172* region was conserved as expected, but there were additional conserved regions next to miRNA/miRNA* (Figure 2A). The Circos analysis of *MIR172a* revealed conserved regions that generate a dsRNA segment of ^∼^15 nt below the miRNA/miRNA* (Figure 2B, 2E, pink line). Furthermore, sequences below this lower stem or above the miRNA/miRNA* tended to be ssRNA in the different species (Figure 2B, 2E, see purple bars). This visualization of the miR172a precursor obtained after our sequence analysis represents the model of the base-to-loop processing of plant miRNAs fairly well, which requires a ^∼^15-17 nt stem below the miRNA/miRNA* that is recognized by a DCL1 complex to produce the first cut (Mateos et al., 2010; Song et al., 2010; Werner et al., 2010; Bologna et al., 2013b; Zhu et al., 2013). MiR393a and miR390a precursors are also known to have a dsRNA region below the miRNA/miRNA* important for their processing (Cuperus et al., 2010; Bologna et al., 2013b). Analysis of their sequences revealed the presence of a conserved ^∼^15-17 stem below the miRNA/miRNA* (Figure 2C, 2F and 2D, 2G). Therefore, the analysis of the *MIRNA* sequences of different species can identify conserved regions that are coincidental with the structural determinants necessary for the precursor processing.

As shown by the Circos-based visualization of miRNA precursors (Figure 2B-G, Supplementary Figure 4 and 5) and the *MIRNA* alignments (Figure 2A, Supplementary 1 and 2), conservation of primary sequences often co-occur with conservation of secondary structures. We also analyzed the existence of compensatory mutations in the *MIRNA* sequence alignments, identifying positions in which the precursor secondary structure is conserved, despite changes in the primary sequence (Supplementary Figure 6). These results are in good agreement with the experimental data showing that the fold-back structure of the precursor is recognized during miRNA biogenesis (Cuperus et al., 2010; Mateos et al., 2010; Song et al., 2010; Werner et al., 2010). The overall analysis generated similar results in dicots alone (Supplementary Figure 4) or dicots and monocots (Supplementary Figure 5), although for a few *MIRNAs*, such as *MIR390a* (Figure 2D and G), there was more divergence in monocots (Figure 2D and G). Therefore, we focused on the analysis of the 30 dicotyledonous species.

**Figure 6:**
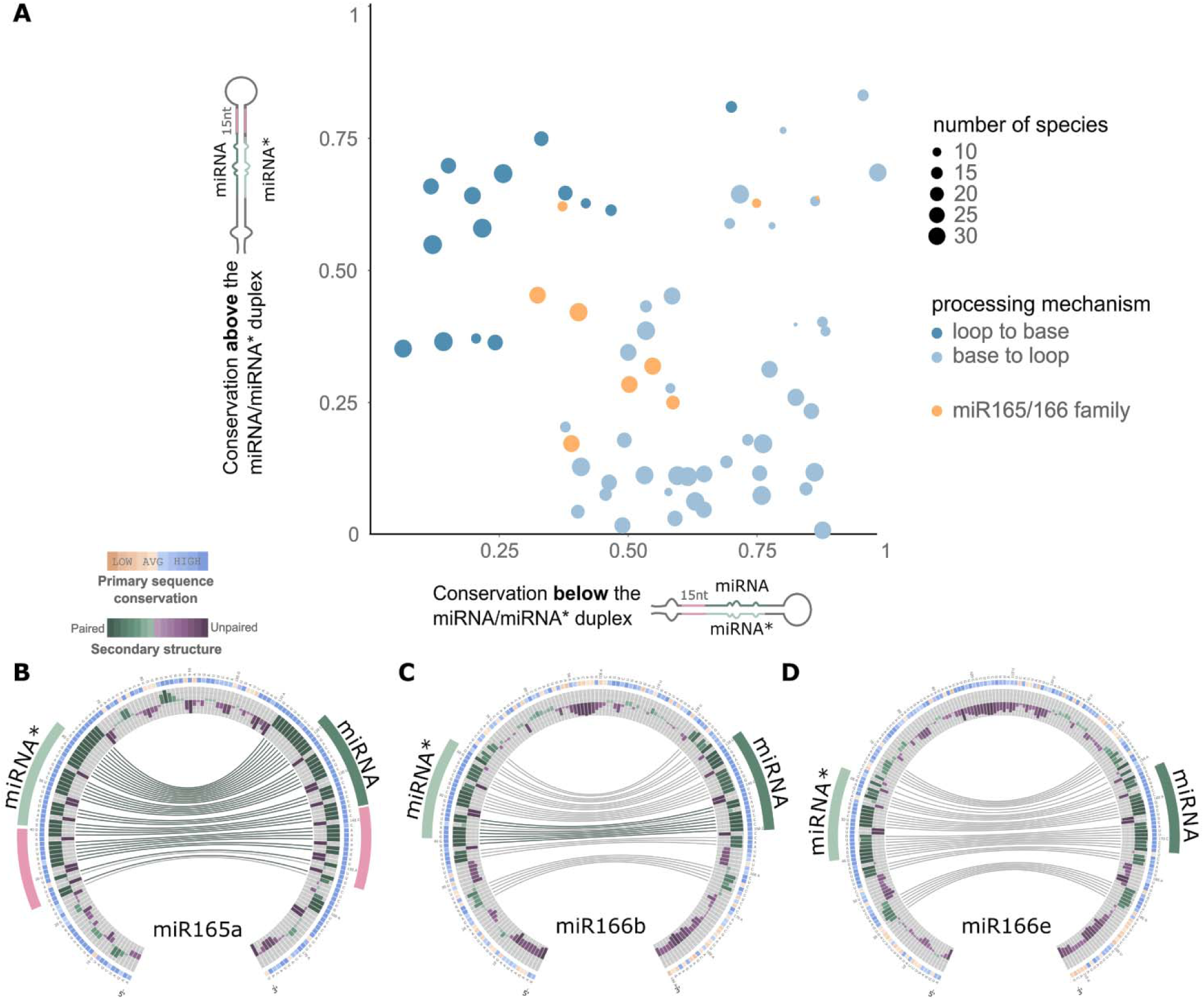
miR166 family members have unusual patterns of conservation. (A) Scatter plot showing mean phastCons conservation score for the sequences above and below the miRNA/miRNA*. We considered 15nt on each arm of the precursor above the miRNA/miRNA*, and 15nt below this region in both arms. The vertical axis represents the average phastCons conservation above the miRNA/miRNA* and the horizontal axis the average phastCons conservation below this region. (B-D) Circos representation of miR165a (B), miR166b (C), and miR166e (D) precursors.

### The precursor sequence conservation is associated to the processing direction

It has been recently shown that several plant miRNA precursors are processed by two DCL1 cuts in a loop-to-base direction (Bologna et al., 2013b). Furthermore, the Arabidopsis miR171a precursor is processed from the base to the loop (Song et al., 2010; Bologna et al., 2013b), while miR171b and miR171c are processed starting from the terminal loop towards the base (Bologna et al., 2013b). We analyzed the sequence conservation of *MIR171a* and *MIR171c* and found strikingly different patterns of conservation (Figure 3A-B). The precursor of miR171a had a conserved dsRNA region below the miRNA/miRNA* region (Figure 3A) like miR172 (Figure 2B). In contrast to the precursor of mi172a and miR171a, in the case of the miR171c there was a conserved region above the miRNA/miRNA*, which determine a dsRNA segment (Figure 3B, pink line). That members of the miR171 family have different patterns of conservation supported our strategy that sought to compare orthologous *MIRNAs* rather than grouping different members of the same family. Other miRNA precursors processed from the loop with two cuts, such as *MIR160a* also have a conserved dsRNA segment above the miRNA/miRNA* (Figure 3C, pink line). Furthermore, the sequence alignments of *MIR160a* (Figure 3D) or *MIR171c* (Supplementary Figure 1 and 2, Supplementary Dataset 2) of different species displayed conserved regions between the miRNA and miRNA*, in contrast to the alignments of *MIR172a* (Figure 2A) or *MIR171a* (Supplementary Figure 1 and 2, Supplementary Dataset 2) that showed additional conserved regions outside the miRNA and miRNA*.

To quantify the degree of sequence conservation, we turned to phastCons (Siepel et al., 2005). We analyzed the conservation in two contiguous regions of 15 nt below the miRNA in the precursor (L1 and L2, L1 being the region adjacent to the miRNA sequence), and one region above the miRNA (U). The regions next to the miRNA* (L1*, L2* and U*) were also analyzed. In base-to-loop precursors, the L1/L1* regions were more conserved than the U/U* (p-value < 1,6e-15, Wilcoxon nonparametric test) and the L2/L2* (p-value < 2,1e-11) (Figure 3E), as expected from the known importance of the dsRNA region immediately below the miRNA/miRNA* for the precursor processing. In contrast, in loop-to-base precursors, the U/U* regions were more conserved than the L1/L1* regions (p-value < 2,4e-07, Wilcoxon nonparametric test) and the L2/L2* (p-value < 1,2e-14) (Figure 3F). Overall, these results confirmed that precursors processed in different directions have specific patterns of sequence conservation.

Next, we analyzed the conservation of young *MIRNAs* present only in Brassicacea species that are processed in a base-to-loop direction. We observed again that L1/L1* were more conserved than L2/L2* (p-value < 3,4e-5, Wilcoxon nonparametric test) and the U/U* (p-value < 0,001) (Figure 3G). In the young *MIRNAs*, however, we did not observe a statistical difference in the conservation of the miRNA/miRNA* and the L1/L1* regions (Figure 3G). Previous analysis of the young *MIR824* in Arabidopsis ecotypes revealed selection of stable precursor sequences (de Meaux et al., 2008). Our results show the importance of the selection of both the miRNA and specific processing determinants during early events of miRNA evolution. However, during a longer period of time, it would be expected that the L region will diversify more than the miRNA/miRNA* duplex.

### miRNAs biogenesis shapes the precursor conservation pattern

The previous analysis was focused on precursors processed by two DCL1 cuts. Plant miRNA precursors can, however, be processed sequentially by three or more cuts (Figure 4A-D, upper panels). The miR319 and miR159 precursors are sequentially processed in a loop to base direction by four DCL1 cuts, which generates additional small RNAs (Addo-Quaye et al., 2009; Bologna et al., 2009; Zhang et al., 2010; Bologna et al., 2013b). The Circos analysis revealed an extended conservation of the secondary structure of these precursors, which generates a dsRNA region of ∼80nt (Figure 4D). This region correlated with the region spanning the four cleavage sites (Figure 4D, green line) and a dsRNA segment above the first cut (Figure 4D, pink line).

Contrasting, the miR394 family and most miR169 family members are processed sequentially by three cuts starting at the base of the precursor (Bologna et al., 2013b). The analysis of these miRNA precursors revealed that they have a conserved dsRNA region of ∼35 nt below the miRNA and miRNA* (Figure 4C, green line, Supplementary Figure 1) that corresponds to the region spanning the first two cuts from DCL1 and a ^∼^15 nt dsRNA stem below the first cleavage site (Figure 4C, pink line). Quantitative analysis using phastCons for these *MIRNAs* revealed that both regions below the miRNA/miRNA* (L1/L1* and L2/L2*) were more conserved than the region above the miRNA/miRNA* (U/U*) (p-value < 1.8e-05, Wilcoxon nonparametric test). In contrast, in the case of MIR319 and *MIR159*, the two contiguous regions above the miRNA/miRNA* (U1/U1* and U2/U2*) were more conserved than the region below (L1/L1*) (p-value < 6.5e-07). Overall, the results show that precursors processed by more than two cuts have concomitantly longer conserved regions than those processed only by two cuts. The extension in the conserved sequence corresponded to an approximately 21 nt dsRNA segment for each additional cut in the processing of the precursor, which is the approximate distance between two DCL1 cuts.

The results show that there is variation in the sequence conservation of plant *MIRNAs*, but that the pattern of sequence conservation can be linked to the processing mechanism of the miRNA precursors (Figure 4A-D). Moreover, this analysis might also be applied to other RNAs or systems. We analyzed the pattern of conservation of animal *MIRNAs* and observed that they have an extended conservation below and above the miRNA/miRNA* (Figure 4E). We think that this conservation might also be linked to the biogenesis of animal miRNAs. While a lower stem below the miRNA/miRNA* is necessary for the first cut by DROSHA (Han et al., 2006), the region above the miRNA/miRNA* might be important for the export from the nucleus to the cytoplasm (Yi et al., 2003; Lund et al., 2004; Zeng and Cullen, 2004).

### Divergence of the precursor length correlates to the processing direction

Next, we analyzed the distance between the miRNA and the miRNA* in the precursors of different species. Analysis of the miR160a precursor, which is processed from the loop to the base revealed that the distance between miR160 and miR160* remained fairly constant in different species with approximately ∼37 nt and ranging from 36 to 40 nt (Figure 5A, right panel). In contrast, the miR172a precursor, which is processed from the base to the loop displayed a larger variability, and the region between miR172a and miR172a* varied from 38 to 114nt (Figure 5A, left panel). A general analysis showed strikingly different patterns of conservation regarding the distance between the miRNA and the miRNA* in different species (Figure 5B). Precursors processed in a base-to-loop direction like miR172a (Cuperus et al., 2010; Mateos et al., 2010; Song et al., 2010; Werner et al., 2010; Bologna et al., 2013b; Zhu et al., 2013) have variable distances between the miRNA and the miRNA* (Figure 5B). In contrast, miRNA precursors experimentally validated to be processed in a loop-to-base direction (Addo-Quaye et al., 2009; Bologna et al., 2009; Bologna et al., 2013b) displayed uniform miRNA-miRNA* distances (Figure 5B, yellow and orange boxes). It has been shown that the terminal region of the precursors processed by two cuts can be largely modified without impairing the miRNA biogenesis (Mateos et al., 2010; Song et al., 2010; Werner et al., 2010), while deletions in the terminal region of precursors processed from the loop significantly affect their processing (Bologna et al., 2009; Bologna et al., 2013a; Kim et al., 2016). We also extended the terminal region of the miR171b and miR319a precursors, and analyzed their importance in vivo. We found that both precursors were not processed after extending their terminal region (Supplementary Figure 7). Overall, the data show that there is an agreement between the conservation of the precursor length and its importance during miRNA processing. We also noted that miR319a was more tolerant to modifications in the precursor length than miR171b and that miR319 precursors were slightly more variable in their length in different species compared to miR171b and miR160 (Figure 5B, Supplementary Figure 7).

**Figure 7:**
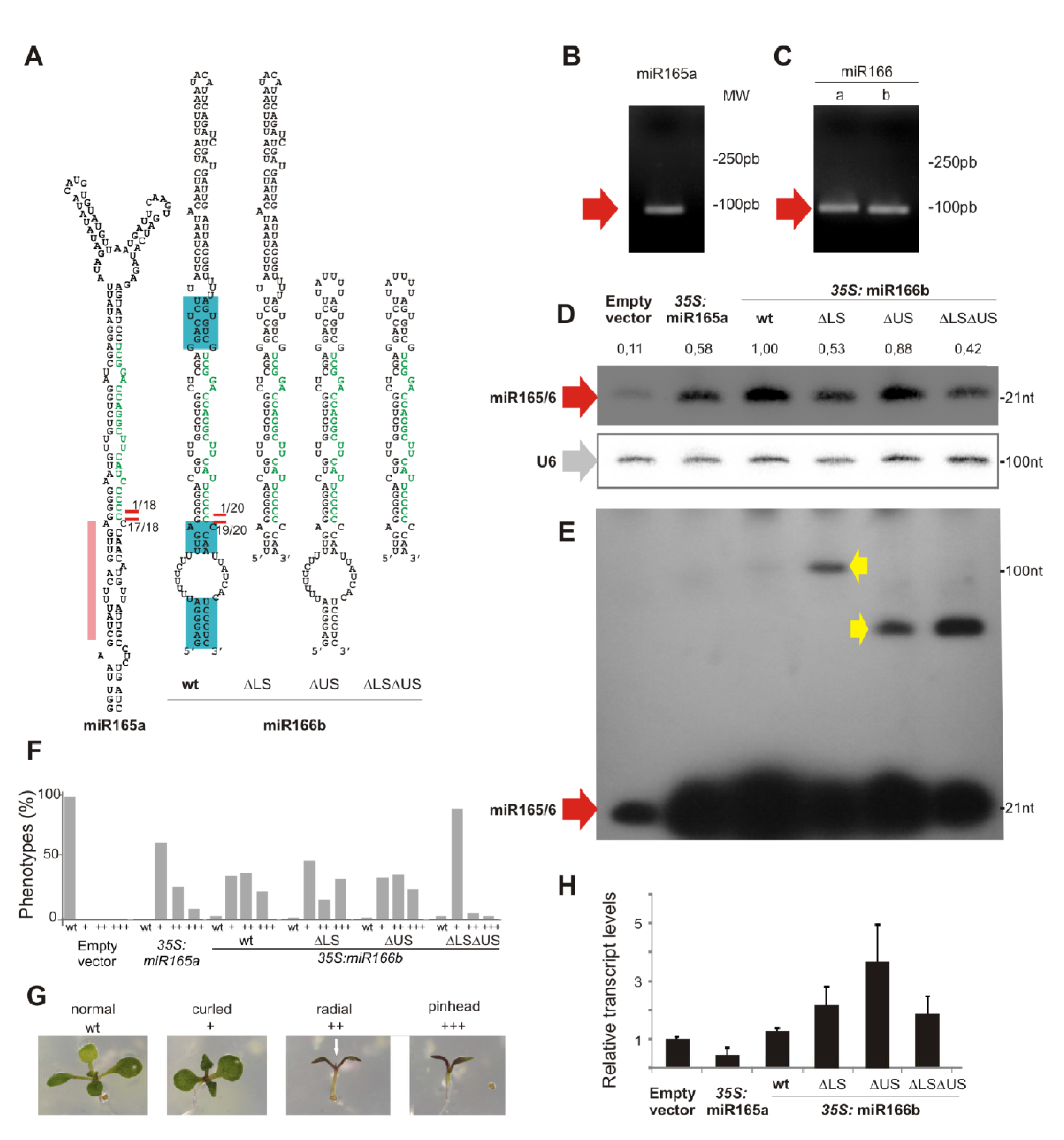
Sequence requirements for miR166 processing in plants. (A) Predicted secondary structures of miR165a, miR166b and mutant miR166b precursors. Red lines and number indicate the number of processing intermediated sequenced in each position. The blue boxes in miR166b precursor indicate the conserved sequences in different species. (B-C) Agarose gels after a modified RACE-PCR to identify processing intermediates of miR165a, miR166a and miR166b precursors. The red arrow indicates the only DNA fragment recovered. (D-E) Small RNA blot of transgenic lines overexpressing miR165a, miR166b and mutant miR166b precursors. At least 20 independent transgenic seedlings were pooled in one sample. The U6 equal loading is shown below. The blot (E) is a longer exposure of (D). Red arrows indicate the miRNA while yellow arrows indicate processing intermediates detected in the blot. (F) Phenotypic defects caused by miR165/miR166 overexpression. Distribution of phenotypic defects in plants overexpressing miR165a, miR166b and mutant miR166b precursors. At least 50 independent primary transgenic plants were analyzed in each case. (G) Photos of the typical developmental defects caused by the over expression of miR165/166. The scale was used to quantify the phenotypic defects on (F). The white arrow indicates a radial leaf-like organ. (H) Pri-miRNA quantification by RT-qPCR of the precursors showed in (D). Error bars indicate the standard error (biological triplicates).

### *MIR166 MIRNAs* have specific patterns of sequence conservation

That the sequence and RNA structure conservation of plant *MIRNAs* correlates with their processing mechanism (Figure 2-4) prompted us to explore whether in certain miRNAs conservation deviates from known processing pathways. We generated a global view of the sequence conservation for all experimentally validated precursors processed from the loop or the base, by comparing the relative conservation above and below each miRNA/miRNA* (Figure 6A). As expected, precursors processed in a base-to-loop direction were more conserved below the miRNA/miRNA*(Figure 6A, light blue dots), while the precursors processed in a loop-to-base fashion were more conserved above the miRNA/miRNA* (Figure 6A, blue dots) (Student’s T-test, P<0.05).

We noticed that members of the miR165/166 family of miRNAs displayed an uneven conservation pattern (Figure 6A, orange dots). The precursor of miR165a, which was more conserved below the miRNA/miRNA* (Figure 6A), showed a conserved ^∼^15 nt stem below the miRNA/miRNA* (Figure 6B), as expected from the base-to-loop processing mechanism. However, other members of the same family such as *MIR166b* and *MIR166e* showed a short conserved stem below the miRNA/miRNA* followed by a large internal loop and a second dsRNA segment (Figure 6C-D). The latter precursors also have a short but conserved stem above the miRNA/miRNA* duplex (Figure 6C-D). The secondary structure of the Arabidopsis miR166 precursors correlated with the conserved structured regions observed in the sequence conservation analysis (Supplementary Figure 4, Figure 7A). These results suggest that different processing mechanisms might account for the biogenesis of miR165/miR166 miRNAs but, most importantly, that several miR166 precursors have conservation patterns that do not match the known processing mechanisms so far described in plants (Figure 4, upper panels).

### Specific sequence requirements for miR166 processing

To study the sequence requirements for the unusually conserved miR165/166 family members, we focused on *MIR166b*. This precursor harbors a conserved dsRNA region of ∼6 nt above the miRNA/miRNA* region, although the structured region in the Arabidopsis precursor is longer (Figure 6C, Figure 7A). Below the miRNA/miRNA*, there is a short stem of ∼4 nt, a large internal loop followed by an additional structured segment (Figure 6B, Figure 7A). Analysis of processing intermediates, revealed a single intermediate for miR166b precursor, as well as for the precursors miR165a and miR166a (Figure 7B-C), which corresponds to a first cut at the base of the precursor (Figure 7A, red lines).

To study the relative importance of the miR166b precursor sequences, we generated a truncated *MIR166b* harboring only the short conserved dsRNA region of 6 nt above the miRNA/miRNA* (Figure 7A, miR166bΔUS). We introduced miR165a, miR166b and miR166bΔUS precursors in Arabidopsis plants under the control of the 35S promoter. Analysis of the primary transcript levels by RT-qPCR revealed that all precursors were expressed in plants (Figure 7H). We detected a higher accumulation of the mature miRNA by small RNA blots from the unusually conserved miR166b precursor than the miR165a. Furthermore, miR166b and miR166bΔUS precursors accumulated similar levels of mature miRNA (Figure 7D), suggesting that a longer dsRNA region above the miRNA/miRNA*, which is not conserved during evolution, is not required for its processing. Then, we deleted the sequences below the conserved short stem of 4nt below the miRNA/miRNA* (Figure 7A, miR166bΔLS). We found that *35S:miR166bΔLS* lines accumulated less small RNA than *35S:miR166b* expressing plants (Figure 7D), indicating that this lower stem had some quantitative effects on the accumulation of the miRNA, but it was not essential. This result was surprising because known precursors processed from base to the loop require a ∼15 nt dsRNA region below the miRNA/miRNA*, and deletions or point mutations in this region completely impair their processing (Cuperus et al., 2010; Mateos et al., 2010; Song et al., 2010; Werner et al., 2010). Previous analysis on the miR172a precursor have shown that loop sequences that are non-essential for miRNA biogenesis can enhance the processing efficiency and have been claimed to stabilize the precursor (Werner et al., 2010), which might also be the case for the deleted lower stem region of miR166b.

Finally, we prepared a mini miR166b precursor leaving only the few conserved bases next to the miR166b/miR166b* (Figure 6C, Figure 7A, miR166bΔLSΔUS). Furthermore, the miR166bΔLSΔUS precursor was expressed and processed in plants producing the mature miRNA, albeit to lower levels than the wild-type precursor (Figure 7D-H). Longer exposure of the small RNA blot for miR166 allowed the detection of processing intermediates, which are consistent with the accumulation of a stem-loop after the first cleavage reaction (Figure 7EC, yellow arrows). These intermediates accumulate at higher levels in the precursors lacking the extended dsRNA regions, suggesting that these stem segments, which are not essential for the miRNA biogenesis, might recruit processing factors that will aid to the biogenesis of miR166b, including the second cleavage reaction. We also scored the phenotypes of primary transgenic plants overexpressing the wild-type and mutant precursors and found a correlation between the mature miRNAs and the developmental defects observed (Figure 7F-G). Most importantly, the mini miR166b precursor was processed *in vivo* and caused developmental defects, confirming that the processing of miR166b does not required a ^∼^15-17 dsRNA region below or above the miRNA/miRNA* as seen in other plant precursors.

## Conclusions

Here, we systematically analyzed *MIRNAs* in different species. We developed a strategy to visualize the conservation of the primary sequence and secondary structure of *MIRNAs*. A general description of plant *MIRNA* sequences revealed regions of sequence conservation that go beyond the miRNA/miRNA*, and that evolutionary footprints can be linked to mechanistic processes occurring during miRNA biogenesis. The approach described here can be used as a practical tool to characterize the constraints of known processing determinants or to provide insights into new mechanisms. Furthermore, the representation allows a quantitative visualization of the conservation of the primary and secondary structures. It is known that single point mutations at specific positions modify the RNA secondary structure and impair the precursor processing (Cuperus et al., 2010; Mateos et al., 2010; Song et al., 2010; Werner et al., 2010), which can explain the conservation at the primary sequence of the structural determinants for miRNA biogenesis.

Extensive biochemical and genetic studies have allowed the characterization of structural determinants that promote the biogenesis of plant miRNAs (Addo-Quaye et al., 2009; Bologna et al., 2009; Cuperus et al., 2010; Mateos et al., 2010; Song et al., 2010; Werner et al., 2010). Our work revealed that the experimentally validated structural determinants can be visualized as clearly conserved regions in the *MIRNAs*. We observed conserved regions corresponding to ∼15-17 bp stems below or above the miRNA/miRNA* depending on the direction of the precursor processing, from the base to the loop or the loop to the base, respectively. Our analysis focused on dicots and monocots, but conserved features corresponding to the lower stem of the miR390 precursor can be found in angiosperms and liverworts (Xia et al., related manuscript submitted to Plant Cell).

The conservation of the miRNA itself during evolution can be explained by its function in the regulation of conserved cognate target sequences [reviewed in (Cui et al., 2016)]. The results presented here show that conservation of the precursor processing mechanism cannot be uncoupled from the miRNA sequence, and that the conservation of structural determinants can be already identified in young *MIRNAs* present only in a group of related species. Studies performed in animals have shown that a group of precursors whose processing is post-transcriptionally regulated require the binding of accessory proteins to their terminal loops, which are conserved during evolution (Michlewski et al., 2008). The approach developed here can be further used to identify and predict mechanistic processes that are specific for a group of miRNA precursors as an alternative to time-consuming experimental approaches.

MiR166 miRNAs fulfill key biological roles including the control of the shoot apical meristem and leaf polarity and overexpression of miR165/166 affects the shoot meristem and leaf polarity [reviewed in (Holt et al., 2014)]. MIR166 precursors have been identified in a wide range of plant species, including mosses (Floyd and Bowman, 2004; Barik et al., 2014). Previous studies have shown that the miRNA/miRNA* duplex of miR165/miR166 have a specific structure that allows the loading into AGO10 (Zhu et al., 2011). The data obtained here suggest that the whole biogenesis of miR166 might have also specific features. That at least some miR166 precursors require only a few bases adjacent to the miRNA/miRNA* region is different to other known miRNAs. The processing of miR166c precursor has been studied in detail *in vitro*, and shown to have a base-to-loop processing mechanism, however in the same system miR166a and miR166b precursors were not processed (Zhu et al., 2013). These results are consistent with different processing mechanisms acting on miR165/miR166 family members. Furthermore, we cannot discard the recruitment of specific cofactors for the processing of these precursors in plants.

The structure of the miR319 precursors is unusual as it has a long foldback with an additional block of sequence conservation below the loop (Palatnik et al., 2003; Axtell and Bartel, 2005; Warthmann et al., 2008; Addo-Quaye et al., 2009; Bologna et al., 2009; Li et al., 2011; Sobkowiak et al., 2012). In this regard, we would like to propose that miR319 and miR166 are two extreme examples: in the long miR319 precursor the structural determinants for its processing are separated from the miRNA/miRNA* region generating an additional block of conservation, while in the miR166b precursor the processing determinants are partially overlapping with the miRNA/miRNA* region.

## Methods

### Identification of plant miRNA precursors orthologous

*MIRNA* sequences belonging to 29 evolutionarily conserved miRNAs present in *A. thaliana* were download from miRBASE release 21. We extended the *MIRNA* sequences to 150 nt outside of the miRNA and miRNA*. Plant genome sequences from 30 dicotyledonous and 6 monocotyledonous species were downloaded from Phytozome (phytozome.jgi.doe.gov, version 11) for the identification of orthologs. We identified putative orthologous genes using a reciprocal Blast hit method using in-house scripts and the NCBI Blast+ package (Altschul et al., 1990). Reciprocal BLAST was done with the *MIRNA* from *A. thaliana* and BLASTing it to the genome of each of the dicotyledonous species. The highest-scoring sequence was taken and BLASTed to the *A. thaliana* database. If this returned the sequence originally used as the highest scorer, then the two sequences were considered putative orthologues. The animal sequences were downloaded from the Ensembl database (Kersey et al., 2016). Homo sapiens *MIRNAs* obtained from miRbase were used to identify orthologous sequences from 13 species: *Bos taurus, Canis familiaris, Equus caballus, Gallus gallus, Gorilla gorilla, Macaca mulatta, Monodelphis domestica, Mus musculus, Ornithorhynchus anatinus, Petromyzon marinus, Sus scrofa and Xenopus tropicalis.*

### Multiple sequence alignments and RNA Secondary Structure analysis

Multiple sequence alignments were performed using command-line version of T-Coffee (version 11.00.8cbe486) (Notredame et al., 2000). We used the slow_pair global pairwise alignment method to build the library, recommended for distantly related sequences. Then we used the +evaluate flag to color the precursor alignments according to its conservation level. Secondary structure prediction from individual precursors in different species were made using RNAfold (Vienna RNA package version 2.1.9) (Lorenz et al., 2011).

### Circos visualization of miRNA precursors

We used Circos (Krzywinski et al., 2009) to make a representation of the miRNA precursors. We put together data from different plant species including multiple alignments with T-coffee, secondary structure with RNAFold and miRNA information. The outer Circos’s data shows the nucleotide sequence of *A. thaliana* precursor, and the color conservation for each position in the consensus of the multiple sequence alignment according to its conservation level output from T-coffee. We have omitted gaps and only the bases within the *A. thaliana* precursor are represented. The inner Circo’s ring shows a histogram of the frequency distribution of paired and unpaired base for that base in the precursor. The degree of conservation of the secondary structure for each miRNA precursor was calculated using structure information in bracket notation from RNAfold. The lines with different colors show the interaction of base pairs in the precursor (green lines mean that these two bases interact in all the analyzed species and the gray lines mean that these two bases interact in at least half of the species).

### Sequence conservation analysis

We used Phast (v1.4) for identifying evolutionarily conserved elements in a multiple alignment, given a phylogenetic tree (Siepel et al., 2005). PhyloFit was used to compute phylogenetic models for conserved and non-conserved regions among species and then gave the model and HMM transition parameters to phastCons to compute base-by-base conservation scores of aligned miRNAs precursors. Using this score, we analyzed the conservation in two contiguous regions of 15 nt below the miRNA in the precursor (L1 and L1), and one region above the miRNA (U). We also considered the cognate regions next to the miRNA* (L1*, L2* and U*). All statistical tests and plots were performed using the R statistical software package. The Wilcoxon test was computed with default parameters and used in PhastCons comparisons between different regions of precursors. For the analysis of miRNAs processed from the base to the loop we used miR164b miR164c, miR165a, miR167a, miR167b, miR167d, miR168a, miR168b, miR169a, miR170, miR171a, miR172a, miR172b, miR172c, miR172d, miR172e, miR390a, miR390b, miR393a, miR393b, miR395a, miR395b, miR395c, miR396a, miR396b, miR397a, miR398b, miR398c, miR399b, miR399c, miR403 and miR827 precursors. For the analysis of miRNAs processed from the loop to the base we used miR156a, miR156b, miR156c, miR156d, miR156e, miR156f, miR156g, miR156h, miR160a, miR160b, miR160c, miR162a, miR162b, miR171b, miR171c precursors. And for the analysis of young miRNAs, we used miR158a, miR158b, miR161, miR771 and miR824 precursors.

### Plant material

All plants used in this work are *Arabidopsis thaliana*, accession Col-0. Plants were grown in continuous light a temperature of 22^<u>°</u>^C. Described phenotypes were scored in at least 50 independent primary transgenic plants.

### Transgenes and precursor analysis

*MIR165a*, *MIR166b*, *MIR319a* and *MIR171b* were obtained from Arabidopsis genomic DNA. Site-directed mutagenesis, plant transformation and scoring of phenotypes were performed as described before (Bologna et al., 2013b), (Zhu et al., 2013). The exact precursor sequences and vectors used here are described in Supplementary Table 1. Cleavage site mapping by modified 5′ RACE PCR was performed as described previously (Bologna et al., 2009), using 10 days old Col-0 seedling. The PCR products were resolved on 3% agarose gels and detected by UV exposure of the ethidium bromide.

### Small-RNA analysis

Seedlings were collected and processed with TRIzol (Invitrogen). Northern blots were performed with 6-12 ug of total RNA resolved on 17% polyacrylamide denaturing gels (7M urea). At least 20 independent transgenic plants were pooled together in one sample. For miR171, miR319 and miR165/166 antisense oligos were 5′end-labelled with [γ-32P] ATP using T4 polynucleotide kinase (Fermentas). Hybridizations were performed as described before (Bologna et al., 2009). The relative miRNA accumulation in the small RNA blots was measured using GelQuant.NET software provided by biochemlabsolutions.com.

Pri-miRNA levels were determined by RT-qPCR. Total RNA (40ng) was treated with RQ1 RNase-free DNase (Promega). The first-strand cDNA synthesis was performed using M-MLV Reverse Transcriptase (Invitrogen). PCR was performed in a Mastercycler ep realplex thermal cycler (Eppendorf) using SYBR Green I to monitor double-stranded DNA synthesis. The relative transcript level was determined for each sample, normalized to the PROTEIN PHOSPHATASE2A cDNA level (Czechowski et al., 2005). For the pri-miRNA we used primer sequences against the CHF3 transcribe regions as described previously (Bologna et al., 2013b).

## Supplemental data

### Supplemental Figures

Supplemental Figure 1. T-Coffee alignments of putative orthologs in dicotyledonous species of the 96 Arabidopsis precursors analyzed in this work.

Supplemental Figure 2. T-Coffee alignments of putative orthologs in dicotyledonous and monocotyledonous species of the 96 Arabidopsis precursors analyzed in this work.

Supplemental Figure 3. RNAfold secondary structure predictions of the miRNA precursors analyzed in this work.

Supplemental Figure 4. Circos-based representation of 96 precursor orthologs in dicotyledonous species.

Supplemental Figure 5. Circos-based representation of 96 precursor orthologs in dicotyledonous and monocotyledonous species.

Supplemental Figure 6. T-Coffee alignments with compensatory mutations.

Supplemental Figure 7. Insertions affect the processing of miR171b and miR319a precursors.

### Supplemental Tables

Supplemental Table 1. List of binary vectors used in this work.

### Supplemental Datasets

Supplemental Dataset 1. Precursor sequences identified and used in this work.

Supplemental Dataset 2. T-coffee alignments.

Supplemental Dataset 3. Information to generate Circos-based visualizations.

## Acknowledgements

We thank Detlef Weigel, Alexis Maizel, Evan Floden, Nicolas Bologna, Carla Schommer and members of the JP lab for discussions and comments on the manuscript. Supported from Bunge and Born and IUBMB Wood-Whelan fellowships to U.C., CONICET fellowships to A.M.L.R, A.L.S, B.M., and J.M.D., J.F.P. is a member of the same institution. Most of the work was supported by grants from Argentinian Ministry of Science, PICT-2012-1780 and PICT-2015-3557, to J.F.P. Supported also by Plan Nacional [BFU2011-28575 to C.N., P.D.]; Center for Genomic Regulation (CRG); ‘Fundació Obra Social la Caixa’ (to E.W.F, M.C); Spanish Ministry of Economy and Competitiveness, ‘Centro de Excelencia Severo Ochoa 2013–2017’ [SEV-2012–0208]; Center for Genomic Regulation (CRG) Funding for open access charge.

